# A grow-and-lock model for the control of flagellum length in trypanosomes

**DOI:** 10.1101/266759

**Authors:** Eloïse Bertiaux, Benjamin Morga, Thierry Blisnick, Brice Rotureau, Philippe Bastin

## Abstract

Several models have been proposed to explain how eukaryotic cells control the length of their cilia and flagella. Here, we investigated this process in the protist *Trypanosoma brucei*, an excellent system for cells with stable cilia like photoreceptors or spermatozoa. We show that the total amount of intraflagellar transport material (IFT, the machinery responsible for flagellum construction) increases during flagellum elongation, consistent with constant delivery of precursors and the previously reported linear growth. Reducing the IFT frequency by RNAi knockdown of the IFT kinesin motors slows down the elongation rate and results in the assembly of shorter flagella. These keep on elongating after cell division but fail to reach the normal length. This failure is neither due to an absence of precursors nor to a morphogenetic control by the cell body. We propose that the flagellum is locked after cell division, preventing further elongation or shortening. This is supported by the fact that subsequent increase in the IFT rate does not lead to further elongation. The distal tip FLAM8 protein was identified as a marker for the locking event. It is initiated prior cell division, leading to an arrest of elongation in the daughter cell. Here, we propose a new model termed grow-and-lock where the flagellum elongates until a locking event takes place in a timely defined manner hence fixing length. Alteration in the growth rate and/or in the timing of the locking event would lead to the formation of flagella of different lengths.

## INTRODUCTION

Cilia and flagella (interchangeable terms) are present at the surface of many eukaryotic cells from protists to humans where they are involved in a range of functions including motility, sensing or morphogenesis. Multiple types of ciliary organisations are encountered from one species to another, and also between different cells in the same organism. Striking variations have been noted in cilia composition, positioning or length, presumably reflecting an optimisation related to the function in a given cell type. The lifespan of cilia is also highly variable, from the transitory existence of some primary cilia to the very stable cilia or flagella of photoreceptors or spermatozoa that show little or no turnover of their microtubules.

Despite extensive variation, each cilium or flagellum exhibits a defined length, a process that has fascinated scientists for decades [1]. To decipher the mechanisms that control length, it is essential to understand how the organelle is constructed. Tubulin is delivered via Intraflagellar Transport (IFT) to the distal end of growing microtubules where incorporation takes place [2-4]. Absence of IFT prevents cilium construction in all organisms investigated so far [5]. The control of flagellum length has mostly been studied in the green algae *Chlamydomonas* [6], a member of the Archeoplastida group. In this organism, flagellar microtubules are highly dynamic and exhibit constant disassembly at their plus end. In such a situation, IFT is essential not only for the construction but also for the maintenance of length [7]. Several models have been proposed to explain the control of length in this context [6]. First, the limiting pool model relies on the production of a fixed amount of flagellar precursors. Once all this material has been used, flagellum assembly is completed. Length would therefore be directly controlled at the level of protein production [8]. Second, the balance-point model proposes that length results from equilibrium between assembly and disassembly rates [7]. At the early phase of assembly, all the IFT material is injected in the flagellum, ensuring a high assembly rate. As the flagellum elongates, the concentration of IFT decreases resulting in a slower assembly rate until it equals the constant disassembly rate. This model assumes that precursor loading is constant at all time and the amount of IFT material injected at the beginning of construction would control the length of the organelle. Third, the cargo-loading model proposes that the amount of tubulin associated to IFT trains controls flagellum length. At the early phase of assembly, a lot of tubulin is imported by IFT in the organelle, ensuring rapid elongation. Later on, the amount of tubulin associated to IFT goes down, resulting in slower elongation [3, 4]. Fourth, morphogenetic control has been proposed in cases where the flagellum is tightly associated to the cell body. Finally, a length sensor could be involved to monitor flagellum length.

In mammals, the mouse sperm flagellum relies on IFT for assembly but not for maintenance, and yet the sperm flagellum does not disassemble after maturation [9]. Another case is the connecting cilium of photoreceptors in the retina. These cells display very low turnover and loss of IFT after assembly only impacts ciliary length after 2-3 weeks, possibly because the cell degenerates [10]. How is length control achieved in the absence of IFT? Spermatozoa or photoreceptors do not lend easily to manipulation. By contrast, protists represent great model organisms, are amenable in the laboratory and several of them exhibit very stable flagella. Here, we selected the protist *Trypanosoma brucei* for investigation for several reasons. First, its axoneme is very stable [11] and relies on IFT for construction [12, 13] but not for length maintenance [14], exactly like in spermatozoa or photoreceptors. IFT remains active after assembly to maintain other elements but not axoneme composition [14]. Second, when trypanosomes infect mammals or tsetse flies, they progress through several stages of development during which they assemble flagella of different length [15-17] and composition [18]. This is reminiscent to what multi-cellular organisms do in different cell types and means that the system is flexible. Third, trypanosomes grow well in culture; they assemble their flagellum in a timely, reproducible and well-characterised manner [19-21]; they are amenable to reverse genetics and IFT has been exhaustively quantified [22]. In the procyclic stage, flagellum replication follows a strict pattern tightly linked to the trypanosome cell cycle (Figure 1A). A new flagellum elongates after duplication of the basal body and grows alongside the existing flagellum [19]. After mitosis, the new flagellum has grown to ~80% of its final length [19, 21] and the cell initiates cytokinesis. One daughter cell inherits the old flagellum and the other one inherits the new flagellum after division. This one continues growing until it reaches the final length of 20 μm [23](Figure 1A).

**Figure 1.**
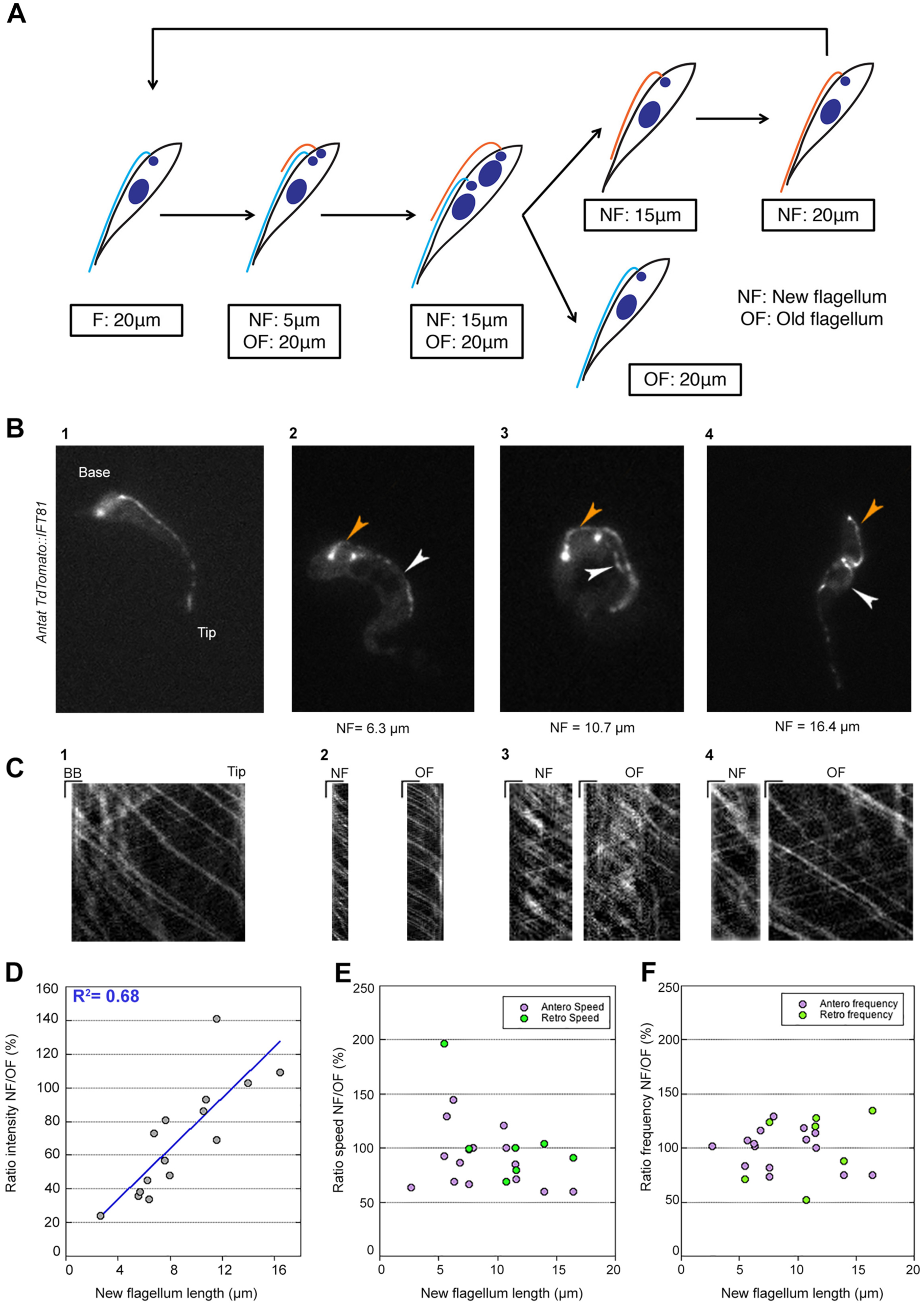
Trypanosome cell cycle and IFT trafficking during flagellum construction. (A) In wild-type conditions, a new flagellum elongates after duplication of the basal body in procyclic trypanosomes. After mitosis, the flagellum has grown to ~80% of its final length [19, 21]. One daughter cell inherits the old flagellum and the other one inherits the new flagellum after division. This one continues growing until it reaches the final length of 20 μm. DNA is shown in dark blue whereas old and new flagella are shown in cyan and orange, respectively. (B) Still images of AnTat1.1E cells expressing a TdTomato::IFT81 from the endogenous locus. Subpanel 1 shows a cell with a single flagellum (Video S1) and subpanels 2-4 show cells at successive stages of flagellum construction (Videos S2, S3 and S4). Orange and white arrowheads indicate the new and the old flagellum, respectively. (C) Kymographs extracted from the corresponding videos where the X axis corresponds to flagellum length (horizontal scale bar, 2 μm) and the Y axis represents the elapsed time (vertical bar, 1s). (D) The ratio between the total TdT::IFT81 fluorescence intensity in the new and the old flagellum was calculated and plotted according to the length of the new flagellum. A linear correlation curve is indicated together with its R^2^ coefficient. (E) The ratio between IFT rates (anterograde transport, magenta circles; retrograde transport, green circles) in the new flagellum and the old flagellum from the same cell was calculated and plotted according to the length of the new flagellum. Retrograde transport is more difficult to detect and data were only incorporated when the signal was sufficiently reliable. (F) Quantification of the ratio between the IFT frequency (anterograde transport, magenta circles; retrograde transport, green circles) in the new flagellum and the old flagellum in the same cell was calculated and plotted versus the length of the new flagellum. See Table S1 for total number of trains analysed. See also Videos S1, S2, S3 and S4.

Here, we investigated flagellum construction in procyclic *T. brucei.* The balance point model is unlikely to apply since IFT is not required for the maintenance of flagellum length in this organism [14]. The tubulin-loading model cannot be tested directly because tagged trypanosome tubulin fails to incorporate into microtubules and does not access the flagellum [24, 25], preventing the visualisation of fluorescent tubulin associated to IFT trains. We show that IFT amounts increase concomitantly with flagellar elongation, in contrast to what has been reported in *Chlamydomonas.* Reducing IFT frequency by IFT kinesin knockdown results in the construction of shorter flagella that keep on elongating after cell division but fail to reach the normal length. This failure is explained neither by a limiting pool of soluble tubulin nor by a control via cell body length. We propose that the flagellum is “locked” after cell division and that flagellum length would hence be fixed at this stage. The locking event is controlled at the cell cycle level, is triggered prior cell division and associated to the addition of a unique marker protein. To integrate these results, we propose a grow-and-lock model where two parameters would dictate the length of the organelle: the growth rate and the timing of the locking event. This model provides an opportunity to explain the control of flagellum length in cells with stable organelles.

## RESULTS

### IFT delivery remains constant during flagellum construction

So far, IFT had only been quantified in mature flagella of *T. brucei* at the procyclic stage maintained in culture [22, 26]. Therefore, IFT activity was examined during flagellum construction and compared to the mature flagellum that remains present during the cell cycle, providing an ideal control [19]. This was carried out in live procyclic trypanosomes that express a fusion protein between the fluorescent Tandem Tomato protein (TdT)[27] and the IFT-B protein IFT81, upon endogenous tagging in the *IFT81* locus [28]. In addition to the bright signal at the base, a succession of motile spots was detected along the length of the flagellum moving in either anterograde or retrograde direction in cells with a single flagellum (Video S1; Figure 1B1) or in those assembling the new one, hence possessing two flagella (Videos S2-4; Figure 1B2-4)(Figure 1). At first glance, IFT behaviour looked quite similar in both growing and mature flagella (Figure 1B). The total amount of fluorescence emitted by the TdT::IFT81 protein in the flagellar compartment was quantified by using the first image of each movie. Plotting the ratio between the total amount of fluorescence in the new flagellum and that in the old one versus the length of the growing flagellum demonstrated a linear correlation between these two parameters (Figure 1D). This data shows that IFT proteins are progressively recruited to the flagellum as it elongates. This means that the IFT amount increases linearly with length hence IFT density per unit of length remains constant during elongation.

Next, kymograph analysis [22] was carried out to quantify IFT rates and frequencies in cells with one (Figure 1B1 and 1C1) or two flagella at different steps of elongation (Figure 1B2-4 and 1C2-4). Kymograph observations revealed brighter individual traces for anterograde transport and less intense traces for retrograde transport as expected [22]. Both IFT speed (Figure 1E) and frequency (Figure 1F) were invariant during flagellum elongation (Table S1). We conclude that the IFT delivery rate remains constant during flagellum construction, which is consistent with the reported linear growth [20, 29].

### Knockdown of IFT kinesins reduces frequency and speed of IFT and results in the assembly of short flagella

In *Chlamydomonas,* reduction in IFT trafficking resulted in the assembly of shorter flagella [7]. To reduce IFT trafficking, we selected to deplete the expression of kinesin II, the IFT anterograde motor. The genome of *T. brucei* encodes two putative kinesin II proteins (Tb927.5.2090 and Tb927.11.13920) but no kinesin-associated protein (KAP) [30, 31]. Individual RNAi silencing of KIN2A or KIN2B did not result in a visible phenotype: cells assembled apparently normal flagella and grew normally in culture (data not shown), suggesting redundancy. Hence simultaneous knockdown of KIN2A and KIN2B was performed following stable transformation of trypanosomes with a plasmid expressing dsRNA of both *KIN2A* and *KIN2B* under the control of tetracycline-inducible promoters [32]. The efficiency of RNAi silencing in *KIN2A2B^RNAi^* cells was confirmed by western blotting using an antibody against KIN2B [31](Figure S1). The signal for KIN2B dropped by at least 8-fold from day 1 and remained low for at least 6 days, confirming the efficiency of RNAi silencing (Figure S1). The frequency and speed of IFT was examined upon transformation of *KIN2A2B^RNAi^* cells with the reporter construct described above allowing endogenous tagging of IFT81 with TdTomato. Trypanosomes were grown in induced or non-induced conditions and IFT was measured in live uniflagellated cells upon kymograph analysis. In control cells, bright anterograde trains were frequently observed, trafficking from the base to the tip of the flagellum where they were transformed to retrograde trains (Figure S2A; Video S5). Kymograph analysis revealed that the average anterograde speed was 1.7±0.5 μm.s^-1^ (n=159 trains from 10 separate cells) and the mean frequency was 0.64 train.s^-1^ (Figure S2B & Table 1). RNAi-induced cells looked different, the signal at the base of the flagellum appeared brighter and adopted a more elongated shape compared to that in control cells (Figure S2C). The train frequency was reduced to 0.37.s^-1^ (n=95) after one day of induction, and down to 0.25.s^-1^ after 4 to 6 days in RNAi conditions (n=125)(Video S6 and Table 1), a phenomenon that is statistically significant. This is visible on the kymograph with fewer traces in induced cells (Figure S2D) compared to control ones (Figure S2B). In addition, IFT trains travelled more slowly when kinesin expression was knocked down: 1.4 μm.s^-1^ at days 4 or 6 instead of 1.9 μm.s^-1^ at day 0 (Figure S2D and Table 1). We conclude that the joint depletion of KIN2A and KIN2B expression reduced IFT delivery by 3-fold in the flagellum.

**Table 1.** IFT anterograde speed and frequency in the *KIN2A2B^RNAi^* cell line in various conditions.

| Conditions | Frequency (train.sec<sup>-1</sup>) | Speed (μm sec<sup>-1</sup>) | Trains |
| --- | --- | --- | --- |
| No tetracycline | 0.63 ± 0.08 | 1.78 ± 0.51 | 159 |
| 1-day induction | 0.38 ± 0.06 | 1.91 ± 0.64 | 95 |
| 4-day induction | 0.28 ± 0.09 | 1.32 ± 0.65 | 62 |
| 6-day induction | 0.25 ± 0.07 | 1.40 ± 0.59 | 63 |
| De-induced (OF) | 0.38 ± 0.11 | - | 161 |
| De-induced (NF) | 0.50 ± 0.13 | - | 239 |

In the case of conventional tetracycline induction of RNAi, only cells possessing a single flagellum was used for the analysis. In the case of the de-induction experiment, only cells with two nuclei and two flagella were measured. IFT speed was not measured in these experiments. OF, old flagella; NF, new flagella. The reduction in frequency is statistically significant in all induced samples compared to the control non-induced ( $p < 0.001$ ). The increase in frequency in de-induced samples is also statistically significant ( $p = 0.01$ ).

Monitoring the culture by microscopy during the course of RNAi indicated the presence of smaller cells with a shorter flagellum (Figure 2A). To quantify this reduction, cells were fixed, processed for immunofluorescence assay (IFA) using the axonemal marker Mab25 and DAPI for DNA staining, and the length of the flagellum was measured. Cells that possessed a single flagellum were first examined. In non-induced samples, the length of the axoneme was on average ~20 μm, as expected [19]. However, the length of the flagellum was shorter during the course of RNAi induction, down to ~9 μm at day 4. Despite the large dispersion, this difference was statistically significant (Figure 2B). Flagellum length remained in that range over the next two days of induction (Figure 2B), and up to 11 days after having triggered kinesin knockdown (not shown). In control non-induced samples, analysis by scanning electron microscopy revealed the typical elongated trypanosome shape with the flagellum attached to the cell body (Figure 2C). By contrast, flagellum length was clearly shorter in induced cells that displayed a shorter cell body (Figure 2D), in agreement with the role of the flagellum in governing trypanosome morphogenesis [12].

**Figure 2.**
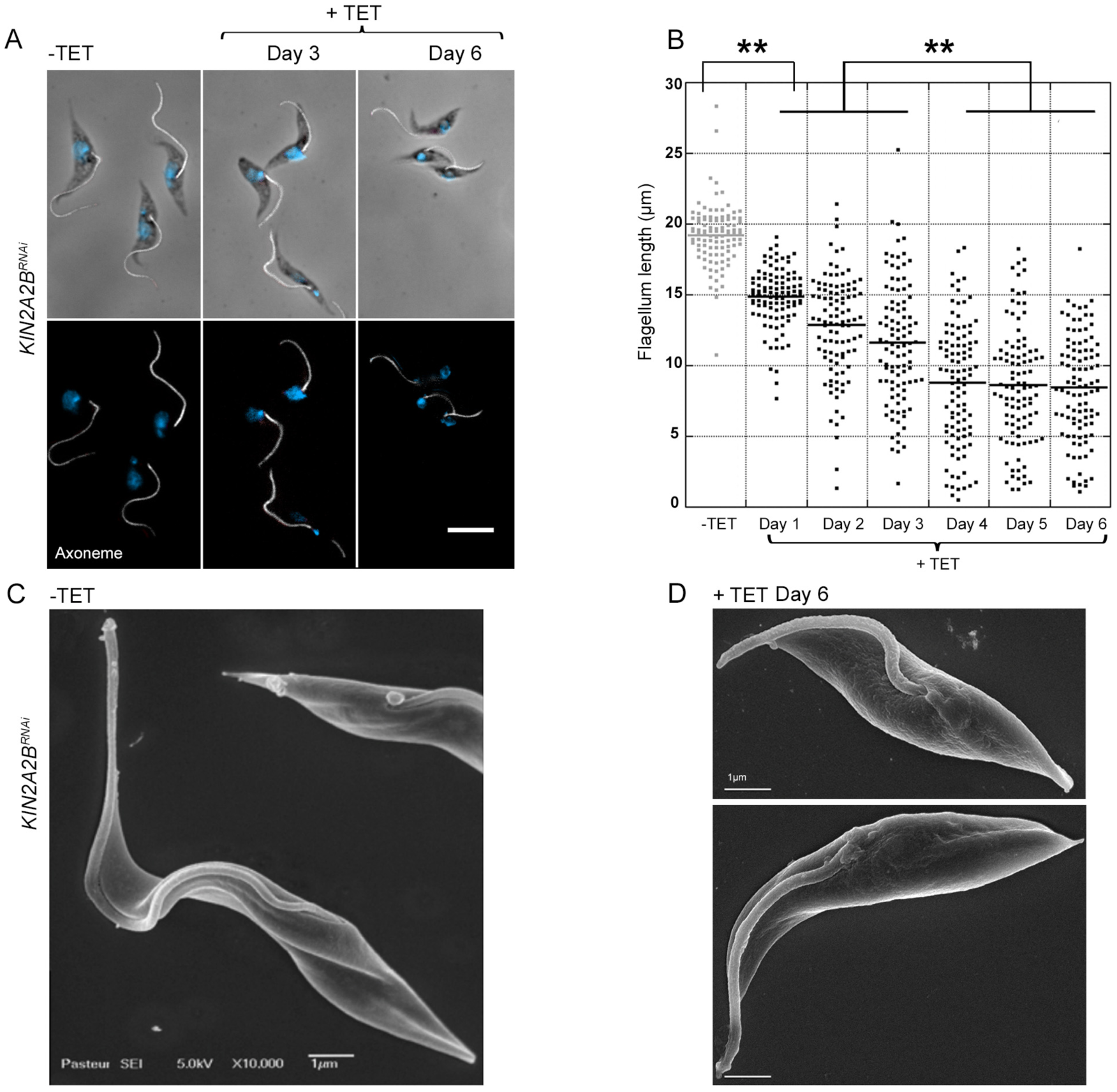
Reduction of IFT train frequency impacts on flagellum length in *KIN2A2B^RNAi^* cells. (A) IFA of non-induced *KIN2A2B^RNAi^* cells, or cells induced for 3 or 6 days as indicated, fixed in methanol, and stained with the Mab25 antibody to detect the axoneme (white). The top panels show the phase-contrast image merged with DAPI (blue) and Mab25 signal (white). Scale bar: 10μm. The bottom panels show Mab25 staining (white) merged with DAPI (blue). (B) Dot plot showing flagellum length during the course of RNAi induction of *KIN2A2B^RNAi^* cells including 100 uni-flagellated cells for each time point. The mean values are indicated with a bold segment. Statistically significant differences are indicated with two stars (p<0.0001). (C-D) Scanning electron microscopy analysis of a non-induced *KIN2A2B^RNAi^* cell showing the typical trypanosome shape and length (C) and of two induced *KIN2A2B^RNAi^* cells for 6 days with shorter flagella (D). See also Figures S1 and S2, and Videos S5 and S6.

The flagellum can be shorter because it is made too short or because it shortens after construction. To discriminate between these two possibilities, trypanosomes at a late stage of the cell cycle (with two nuclei [19]) were investigated. In control non-induced cells, the length of the new flagellum was ~15 μm whereas that of the old flagellum was ~20 μm (Figure 3A, grey symbols on Figure 3B) as expected because construction is not completed before cell division [19]. This was confirmed by scanning electron microscopy of dividing cells: when the cleavage furrow was visible, the length of the new flagellum was shorter than that of the old one (Figure 3C). In induced *KIN2A2B^RNAi^* cells, the average length of the new flagellum was around 7 μm, for all induction times examined and that difference was statistically significant (Figure 3A, black symbols on Figure 3B). Analysis by scanning electron microscopy revealed that the new flagellum is much shorter and that the daughter cell is smaller (Figure 3D) compared to the non-induced cells (Figure 3C). This implies that the new flagellum is made too short when the cell is about to divide. Examination of the same cells showed that the old flagellum was almost twice longer than the new flagellum (12 μm, black symbols, Figure 3B). This indicates that the short flagella continue to grow after cell division but fail to reach their normal length (20 μm). A summary of flagellum formation in KIN2A2BRNAi cells is presented at Figure 3E. At least two main hypotheses directly related to the presence of a shorter cell body can be proposed to explain this result. Flagellum elongation could be restricted (1) by a limited pool of soluble tubulin to sustain further elongation [8] or (2) by the presence of a short cell body to which the flagellum is anchored.

**Figure 3.**
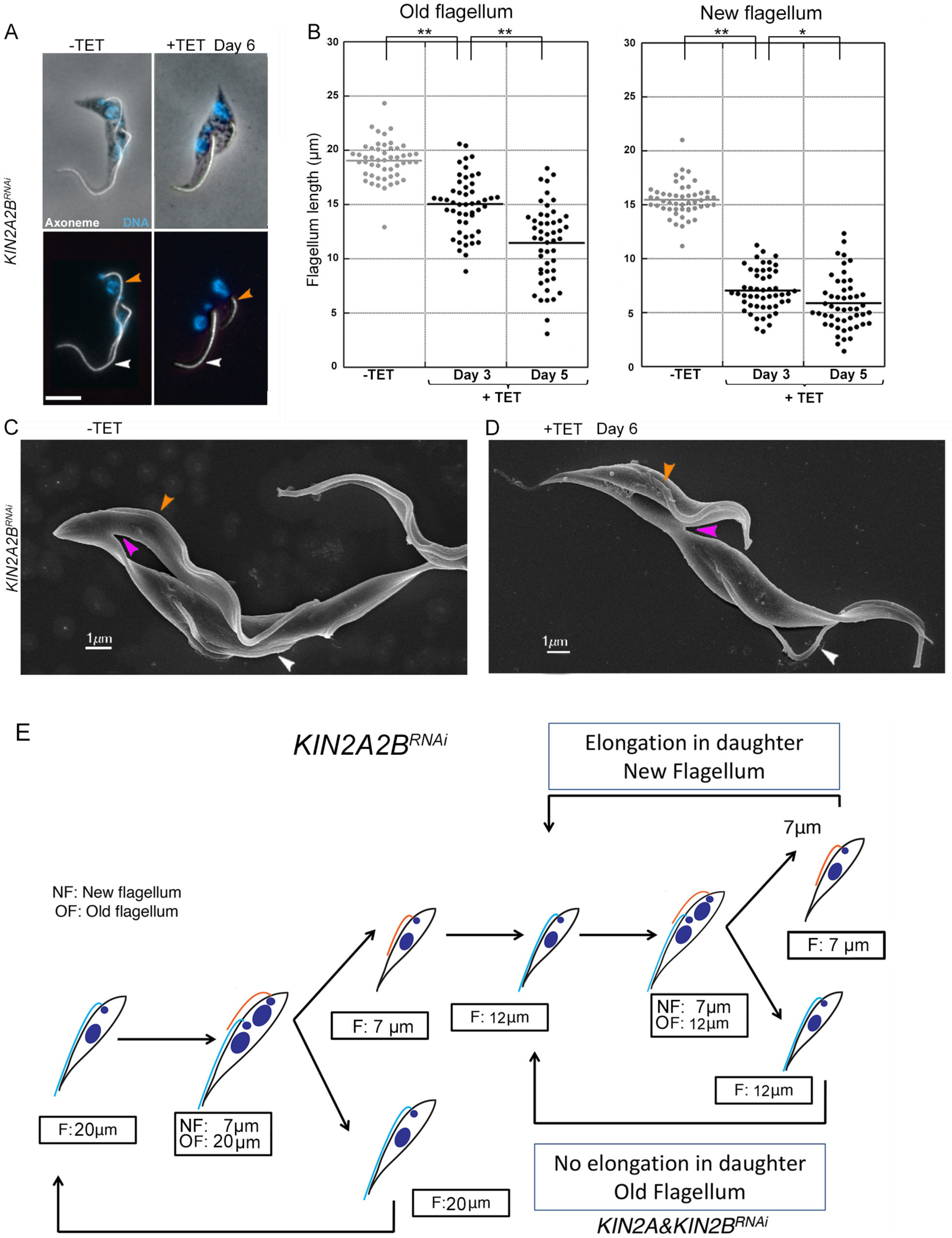
The new flagellum is built shorter upon reduction of IFT train frequency in *KIN2A2B^RNAi^* cells. (A) IFA images of non-induced and 6-day induced *KIN2A2B^RNAi^* cells obtained after methanol fixation and staining with the Mab25 antibody labeling the axoneme (white). The top panels show the phase-contrast images merged with DAPI (cyan) and the Mab25 axonemal marker (red) and the bottom ones show the Mab25 signal (white) merged with DAPI (cyan). Scale bar: 10μm. (B) Dot plot showing the length of old and new flagella during the course of RNAi induction of *KIN2A2B^RNAi^* measured in cells possessing 2 kinetoplasts and 2 nuclei (n= 50 cells for each time point). The mean values are indicated with a bold segment. Statistically significant differences are indicated with one (p<0.001) or two stars (p<0.0001). (C) Scanning electron microscopy pictures of non-induced (C) and induced *KIN2A2B^RNAi^* cells after 6 days (D). The purple arrow indicates the cleavage furrow. Orange and white arrowheads show the new and the old flagellum, respectively. (E) Cartoons showing that the new flagellum only reaches 7 μm at the time of cell division in *KIN2A2B^RNAi^* cells but elongates up to 12 μm until it matures in the next cell cycle. DNA is shown in dark blue whereas old and new flagella are shown in cyan and orange, respectively.

### The limited cytoplasmic pool model or the morphogenetic model do not explain the failure of *KIN2A2B^RNAi^* cells to elongate their flagella

To challenge a regulation by the pool of precursors, the amount of soluble tubulin was determined using cell fractionation in detergent to separate a cytoskeletal and a soluble fraction [33]. Analysis of non-induced *KIN2A2B^RNAi^* cells demonstrated the existence of a low-abundance pool of soluble tubulin (Figure S3) in agreement with previous studies [34]. A similar (possibly even more abundant) pool of soluble tubulin was found in *KIN2A2B^RNAi^* cells after RNAi knockdown (Figure S3). We conclude that the amount of soluble tubulin is not the limiting factor that would cause flagella to be shorter in *KIN2A2B^RNAi^* cells after RNAi knockdown.

The *T. brucei* flagellum is attached to the cell body via a sophisticated cytoskeletal network [35] and this could provide an original way to control flagellum length. This hypothesis is somehow contradicted by the fact that mutants with defects in the adhesion machinery assemble detached flagella of apparently normal length [36-38]. However, these flagella are completely detached from the cell body with the exception of the anchoring via the flagellar pocket and might be regulated differently compared to *KIN2A2B^RNAi^* cells where the short flagella are properly attached to the cell body. To challenge this possibility, a “de-induction” experiment was attempted where *KIN2A2B^RNAi^* cells were grown in RNAi conditions for 6 days, resulting in the formation of short flagella as observed above, followed by tetracycline wash out, leading to expression of fresh KIN2A and KIN2B and restoring IFT (Figure 4A). At the time of tetracycline wash out, cells exhibit a short cell body. If this were a limiting factor, it should prevent further elongation of the new flagellum since it is formed on a short cell body. The return of IFT trafficking in old and new flagella was monitored in live cells by the presence of a fusion protein between mNeonGreen [39] and IFT81 expressed from the endogenous *IFT81* locus [40]. After 16 hours without tetracycline, the frequency of IFT is increased by ~2-fold in both the new and the mature flagellum (Table 1). This was confirmed by immunofluorescence staining with an anti-IFT172 antibody that revealed close to normal signal in both flagella (Figure 4B). This increase in IFT frequency is accompanied by an increase of the length of the new flagellum (Figure 3) that reaches 11.7 ± 3.9 μm (grey circles) instead of 8.25 ± 3.1 μm (black circles). This result contradicts the morphogenetic model for flagellum length control. Furthermore, examining the ratio between the length of the new and the old flagellum revealed further interesting observations (Figure 4D, grey circles). In close to one third of de-induced cells (16/55), the new flagellum was longer than the mature one; something that was never observed in induced *KIN2A2B^RNAi^* cells (Figure 4D, black circles) or in control cells [21]. Several of these cells grew flagella that extended well beyond the cell body (Figure S4). We conclude that flagellum length is not controlled by the size of the cell body.

**Figure 4.**
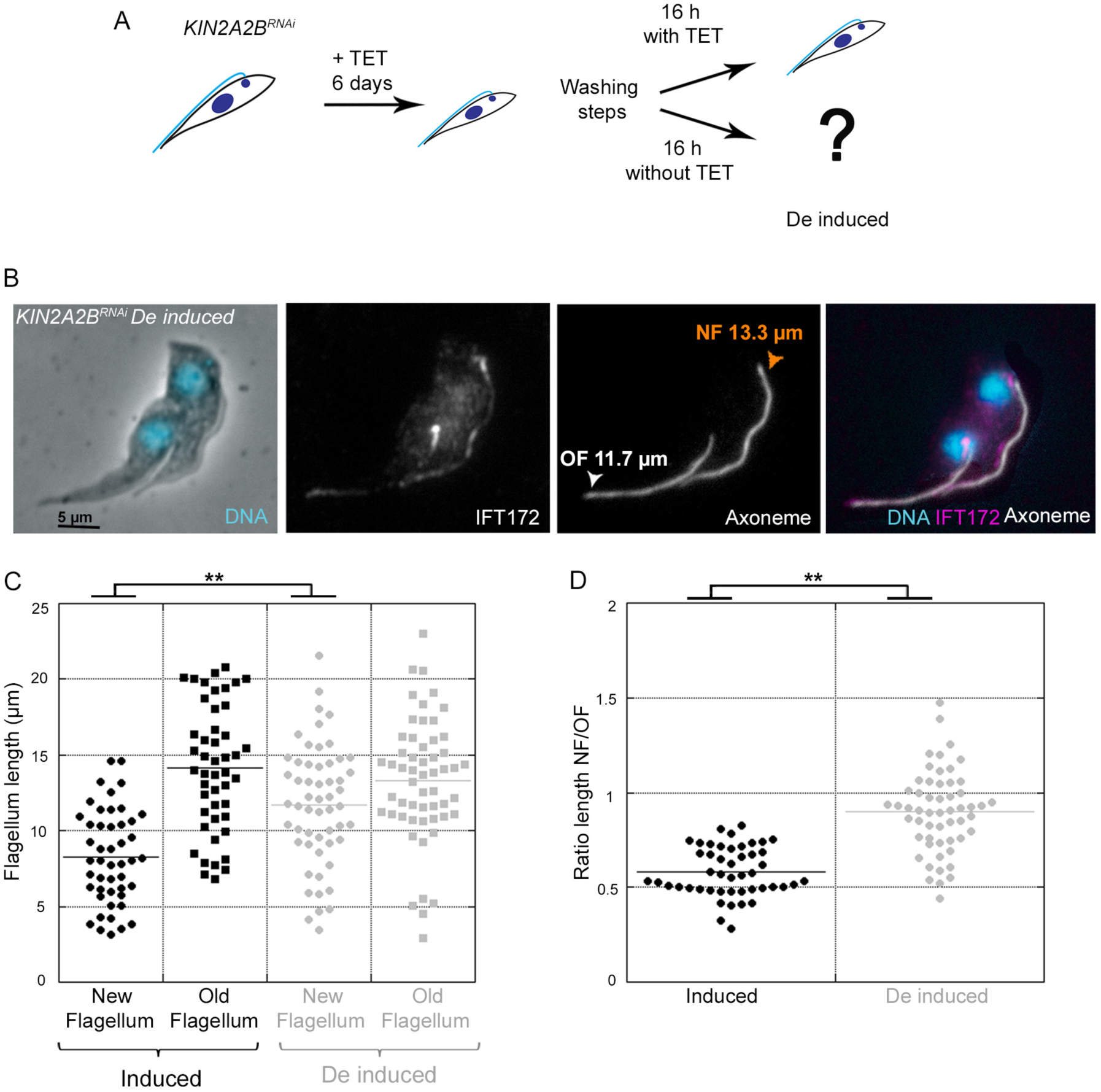
De-induction of RNAi leads to an increase of IFT trafficking in *KIN2A2B^RNAi^* cells but has no impact on the length of the mature flagellum. (A) Schematic representation of the de-induction experiment. *KIN2A2B^RNAi^* cells were grown in RNAi conditions for 6 days before extensive washings to remove tetracycline and returned to culture with (control) or without tetracycline (de-induced) for 16 hours. (B) IFA pictures of a de-induced *KIN2A2B^RNAi^* cell undergoing cytokinesis, stained with the anti-IFT172 antibody and with the Mab25 antibody targeting the axoneme as indicated. The first image shows phase contrast and DAPI staining (cyan). The merged panel contains DAPI (cyan), Mab25 (white) and IFT172 (magenta) signals. Scale bar: 5μm. (C) Dot plot showing the length of old and new flagella in *KIN2A2B^RNAi^* cells that were grown in RNAi conditions (left, black symbols) and in de-induced cells for 16 hours (grey symbols). This was measured in cells possessing 2 kinetoplasts and 2 nuclei (n=54 cells for induced samples and n=49 for de-induced samples). (D) Dot plot representing the ratio between the length of the new flagellum and that of the old flagellum. The mean values are indicated with a bold segment. Statistically significant differences are indicated with two stars (p<0.0001). See also Figure S4.

### Increase in IFT in mature flagella does not result in modification of their length

The “de-induction experiment” also allows to evaluating the length of the old flagellum once IFT has been restored close to normal values. In induced conditions, the average length of the old flagellum was 14.2 ± 4.1 μm (black squares, Figure 4C), as previously observed (Figure 3B). Strikingly, the length of the old flagellum remained unchanged after de-induction at 13.9 ± 4.08 μm (grey squares, Figure 4C). This demonstrates that a 2-fold increase in IFT cannot rescue flagellum length after maturation. This further proves that the balance point model based on different concentrations of IFT material affecting the equilibrium between assembly and disassembly does not apply to this type of flagellum. Indeed, this model would predict an elongation of the flagellum upon increase of the IFT frequency [41]. It also means that something else should prevent flagellum elongation, knowing that soluble tubulin is available and that the cell body is not a restricting factor.

### A locking event is initiated prior cell division

We propose that a locking event of unknown nature could take place once the flagellum matures and would prevent further elongation or shortening. What could control the timing of this event? Since flagellum assembly is intimately linked with the progression through the cell cycle in trypanosomes as in other protists [19, 42, 43], we propose that this putative locking event could be controlled by a cell cycle-dependent mechanism rather than at the flagellum level. When a procyclic trypanosome divides, the new flagellum has reached ~80% of the length of the old flagellum meaning that elongation continues up to 20 μm in the daughter cell inheriting that flagellum. Post-division elongation has been experimentally proven but not quantified so far [23] and the events leading to its arrest are unknown. An arrest of flagellum growth could occur when the flagellum becomes mature after cell division or could be triggered by a specific signal prior cell division hence becoming effective in the daughter cell.

To discriminate these two possibilities, cell division was inhibited. If the locking event takes place after division, it should be possible to restore normal flagellum length since flagellum growth should continue unabated. If it were initiated before cell division, only limited growth would be possible after activation of the signal. *KIN2A2B^RNAi^* cells were grown in the presence of 10 mM teniposide, a drug that interferes with mitochondrial DNA segregation but neither with basal body duplication nor with flagellum elongation [44]. This resulted in the expected arrest of cell division that is clearly visible on the growth curve (Figure S5). *KIN2A2B^RNAi^* cells induced for 5 days were incubated in the presence of 10 mM teniposide. After incubation, cells were fixed and processed for IFA with the axonemal marker Mab25 whilst DNA was stained with DAPI. In controls without teniposide, the mitochondrial DNA segregated normally and cells progressed to the typical pattern with 2 kinetoplasts and 2 nuclei preceding cell division (Figure 5A, top panels). By contrast, kinetoplasts failed to fully segregate in the presence of teniposide (Figure 5A, bottom panels, white arrow), inhibiting cell division. In the absence of teniposide, the ratio between the length of the new and the old flagellum in induced *KIN2A2B^RNAi^* cells was about 60% (Figure 5B, light grey), in agreement with previous measurements (Figure 3B). Remarkably, after 16 hours of incubation with teniposide, the ratio was close to 100% (Figure 5B, dark grey, left bars), meaning that the length of the new flagellum had reached the length of the old flagellum at ~12 μm but still had failed to reach the normal 20 μm. However, this short length could be due to slow elongation because of the reduced IFT frequency and not to a locking event. *KIN2A2B^RNAi^* cells were therefore incubated for 24 hours in the presence of teniposide. These extra 8 hours in teniposide did not result in an increase of new flagellum length that remained stuck at 12 μm (Figure 5B, dark grey, right bars). We conclude that a signal is triggered prior cell division and impacts elongation definitely.

**Figure 5.**
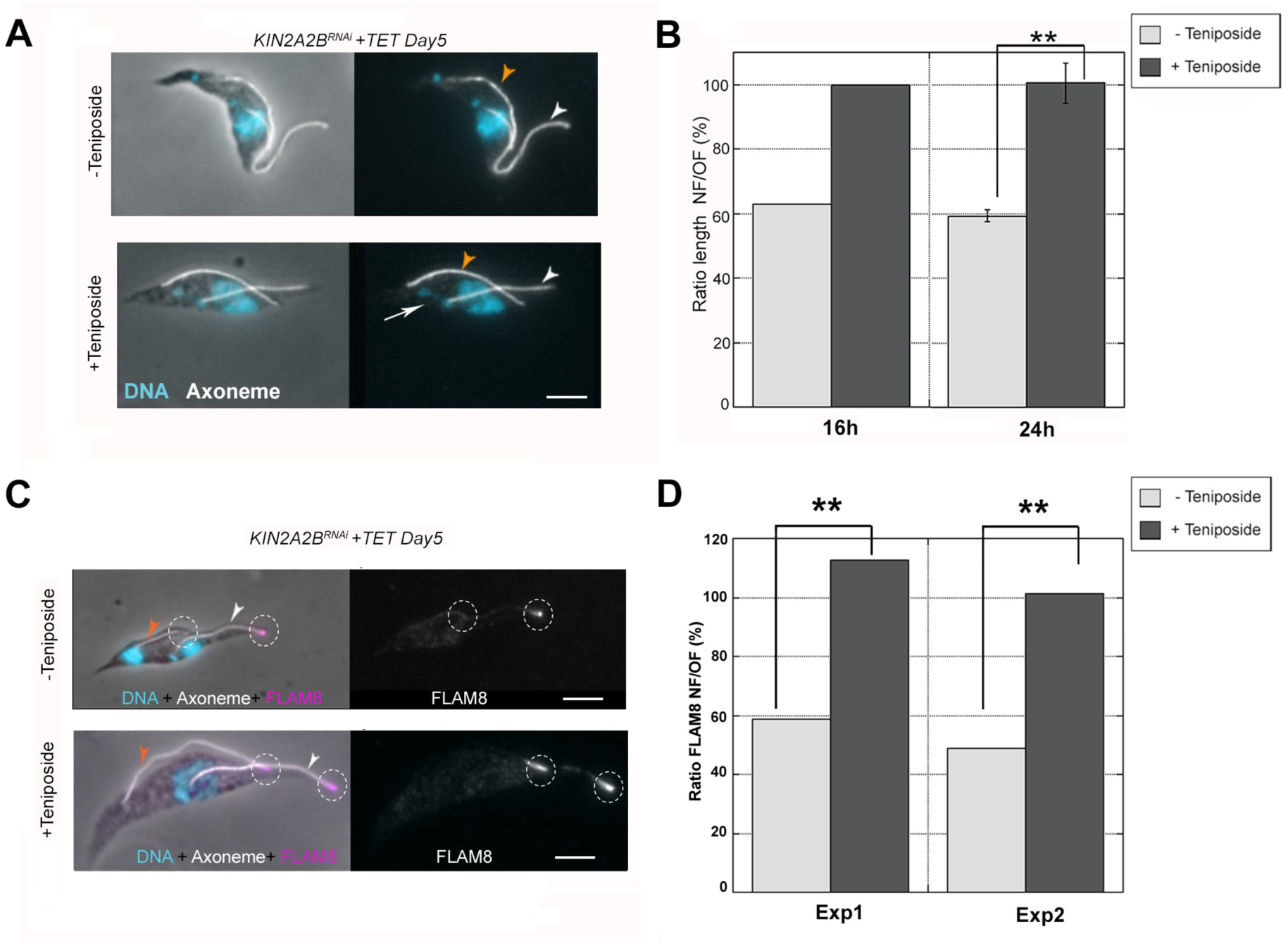
The event leading to flagellum maturation is triggered prior cell division. (A) IFA pictures of 6-day induced *KIN2A2B^RNAi^* cells that were left untreated (top panels) or treated for 24 hours with teniposide (bottom panels), stained with the Mab25 antibody targeting the axoneme (white) and DAPI labeling DNA (cyan). The left panels show the phase-contrast image merged with DAPI (cyan) and Mab25 signal (white). The right panels show the Mab25 signal (white) and DAPI (cyan). Orange and white arrowheads show the new and the old flagellum, respectively. The white arrow shows the bridge linking the kinetoplasts after treatment with teniposide. Scale bar: 5μm. (B) Ratio between the length of the new flagellum and the old flagellum for 6-day induced *KIN2A2B^RNAi^* cells treated (dark bars) or not (white bars) with teniposide during 16 hours (left, one experiment) or 24 hours (right, three independent experiments with standard deviation). For the 16 hour-experiment, n=40 cells for teniposide-treated cells and n=62 cells for the untreated control. For the 24 hour-experiment, n=112 cells for teniposide-treated cells and n=163 cells for the untreated control. Statistically significant differences are indicated with two stars (p<0.0001). (C) IFA pictures of 6 day-induced *KIN2A2B^RNAi^* cells non-treated or treated for 24 hours with teniposide, fixed in methanol and stained using the Mab25 antibody to detect the axoneme (white), the anti-FLAM8 (magenta) and DAPI (cyan). The left panels show the phase-contrast image merged with DAPI (cyan), the anti-FLAM8 (magenta) and Mab25 antibody (white). The right panels show the anti-FLAM8 signal only (white). Orange and white arrowheads show the new and the old flagellum, respectively. The white circles are centered on the FLAM8 signal. Scale bar: 5μm. (D) Ratios between the FLAM8 fluorescent signal intensity in the new and the old flagellum in 6 day-induced *KIN2AB^RNAi^* cells treated (n=109 cells) or not (n=60 cells) with teniposide during 24 hours. Two independent experiments are shown. Statistically significant differences are indicated with two stars (p<0.0001). See also Figure S5 and S6.

One could consider that *KIN2A2B^RNAi^* cells behave differently than wild-type cells for various reasons (shorter flagellum length, shorter cell size, reduced motility) and this might not reflect the normal situation. Therefore, we analysed the impact of an inhibition of cell division in wild-type cells. Blocking cell division should result in an increase of the new flagellum from 80% (length of the new flagellum in wild-type cells) to 100% of the length of the old flagellum. In the absence of teniposide, the ratio between new and old flagella in cells about to divide was close to 80% as expected [21, 45](Figure S6A, top panels and S6B, light grey bars). After incubation with teniposide, the length of the new flagellum reached that of the old flagellum but did not elongate further (Figure S6A, bottom panels and S6B, dark grey bars). We conclude that when cytokinesis is blocked upon inhibition of mitochondrial DNA segregation, the signal that results in locking flagellum length is triggered prior cell division.

### A molecular marker to assess flagellum maturation

In the case of a locking event inhibiting flagellum elongation or shortening, a modification of this flagellum might be expected. The data above indicate this must be progressive since it is initiated prior cell division and leads to an elongation arrest only after division. We therefore searched for candidate molecules that accumulate towards the late phase of flagellum elongation and could serve as markers for flagellum maturation. The FLAM8 protein appeared as an attractive candidate. This large protein (3,075 amino acids) of unknown function was discovered in a proteomic study of purified trypanosome flagella [45]. It is abundant at the distal tip of mature flagella and detected in very low concentrations in short new flagella. However, its amount increases during elongation to reach 40% of that of the old flagellum just prior cell division [45]. It means that a significant increase must happen after cell division, which is compatible with the findings above.

If FLAM8 is indeed a marker of flagellum maturation, it should accumulate in the new flagellum of teniposide-treated cells. Therefore, cell division was inhibited using teniposide in induced *KIN2A2B^RNAi^* cells exactly as above. Cells were fixed and stained by IFA with an anti-FLAM8 antibody, the Mab25 antibody as an axonemal marker, and DAPI to label DNA. When induced *KIN2A2B^RNAi^* cells were not treated with teniposide, FLAM8 was abundant at the tip of the old flagellum, but was present in very low amounts or below detection level in the new flagellum (Figure 5C, top panels). By contrast, in cells treated with teniposide for 24 hours, the new flagellum had elongated further as described above and the FLAM8 signal at its tip was much brighter and looked similar to that at the tip of the mature flagellum (Figure 5C, bottom panels). A region of interest was defined around the tip of the new and the old flagella and the total amount of fluorescence was quantified. In untreated induced *KIN2A2B^RNAi^* cells at an advanced step of their cell cycle, the ratio of FLAM8 signal intensities between the new and the old flagellum was ~60% (Figure 5D, light grey). However, for induced cells treated with teniposide (where the new flagellum reached the length of the old flagellum), the FLAM8 ratio increased to 100% (Figure 5D, dark grey). This difference was statistically significant. We conclude that the molecular composition of growing and mature flagella is different and that a maximum amount of FLAM8 marker is found in flagella that do not elongate further.

## DISCUSSION

Multiple models have been proposed to explain the control of the length of cilia and flagella [6, 8]. However, the new data reported here indicate that most of the existing models fail to explain how trypanosomes proceed. First, the presence of a soluble pool of tubulin of apparently equivalent abundance in control cells and in induced *KIN2A2B^RNAi^* trypanosomes that exhibit short flagella does not support the model based on a limited pool of components [8]. Second, the ability of *KIN2A2B^RNAi^* cells to assemble fairly long new flagella that grow beyond the short cell body demonstrates that a morphogenetic control via the cell body and the cytoskeletal attachment zone system [35] is also unlikely. Third, although the balance point model did not appear relevant from the beginning since flagellum length is maintained normally in the absence of IFT [14], it is formally ruled out here by the fact that the short mature flagellum does not elongate despite a 2-fold augmentation of IFT trafficking in de-induced *KIN2A2B^RNAi^* cells, in contrast to what would be predicted [41]. Finally, the cargo-loading model could not be challenged directly because tagged trypanosome tubulin fails to enter flagella and to be incorporated in flagellar microtubules [24, 25].

One marking feature of several experiments reported here is the fact that the old flagellum does neither elongate nor shorten, whatever the attempted manipulation (modification of IFT frequency, inhibition of cell division). Thus, we propose that this flagellum is locked. This could be ensured by the addition of a cap at the distal end of microtubules, the stabilisation of tubulin by post-translational modification or by preventing fresh tubulin to access the flagellum. First, the addition of a cap at the tip of microtubule doublets could prevent further assembly or disassembly. The identification of the distal tip FLAM8 protein as a marker of maturation goes along this line but it does not necessarily mean that this protein inhibits elongation. So far, a cap structure has not been detected on electron micrographs of *T. brucei* procyclic flagella [46] but the morphology of the axoneme tip looks very different between growing and mature flagella: whilst it looks disorganised in elongating flagella with some microtubule doublets very close to the central pair and others further away and in contact with the membrane, it is nicely and regularly structured in mature ones [47]. A cap could protect against the action of the depolymerising kinesin 13 that has been connected to flagellum length control in the related protists *Leishmania major* [48] or *Giardia intestinalis* [49] but whose contribution appears minor in *T. brucei* [50]. A second possibility is the level of some post-translational modifications of tubulin that alter the dynamics of microtubules. The tip of the growing flagellum contains a large amount of tyrosinated tubulin that is not detected in the mature flagellum [51]. Exhaustive detyrosination could protect the axoneme from disassembly. The locking event could take place at the base of the flagellum, by preventing access of fresh tubulin to the mature organelle.

The locking of the flagellum leads to the consideration of a new model for the control of flagellum length that we termed the “grow-and-lock” model. It proposes that the flagellum elongates to a stage where a signal blocks further elongation or shortening by inducing a modification that locks the structure in a stable (or mature) configuration (Figure 6A). A cell can produce a shorter flagellum either by a slower growth rate (Figure 6A, magenta curve) or by an earlier initiation of the locking event (Figure 6B, magenta). Conversely, a longer flagellum can be constructed using a faster growth rate (Figure 6A, green curve) or by delaying the timing of maturation (Figure 6B, green). Both parameters could be shifted together to achieve the desired length. Experiments performed on *KIN2A2B^RNAi^* cells provide interesting support for the model. First, a reduction in IFT frequency leading to a slower growth rate results in the formation of a shorter flagellum, as predicted by the model (Figure 6A). This is reverted upon an increase of IFT frequency in the new flagellum but not in the old one, consistent with it being locked. Second, blocking cell division allows the flagellum to elongate further, again in agreement with the predictions about the timing of the locking event (Figure 6B).

**Figure 6.**
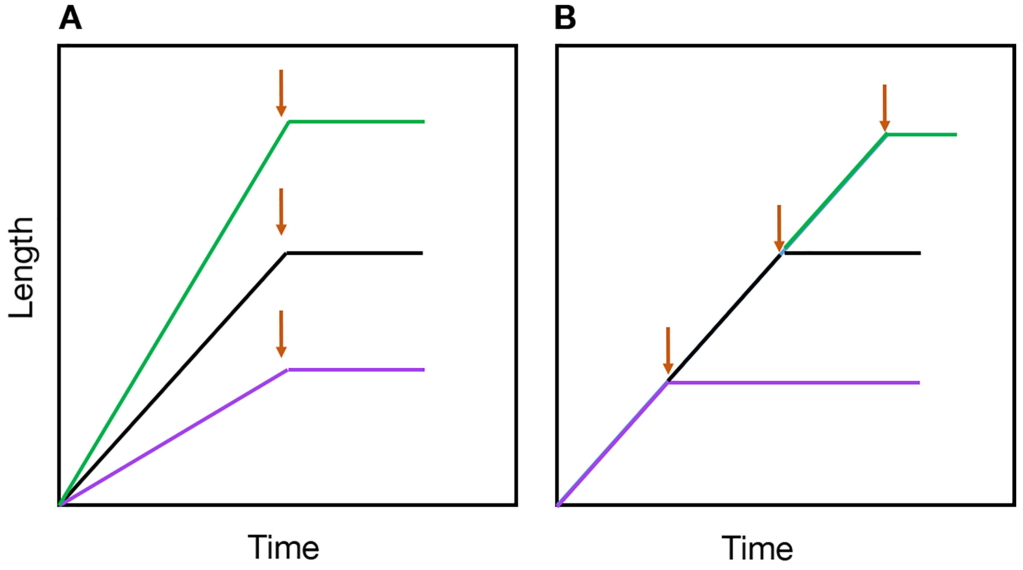
The grow-and-lock model to explain the control of flagellum length in *T. brucei*. (A) A slower (magenta) or faster (green) growth rate allows the production of shorter or longer flagella without having to change the timing of the locking event. (B) The flagellum elongates with a linear growth rate until a point where it is locked (arrow) and shows neither assembly nor disassembly. Keeping the same growth rate but triggering the locking event earlier (magenta) or later (green) will result in the formation of shorter or longer flagella.

The grow-and-lock model has potential for cilia and flagella that do not disassemble their microtubules after assembly and maturation. Locking the length of the axoneme in a mature state could have significant advantages for cellular functions. Protein turnover costs energy and this would dispense the cell of a potentially costly maintenance process. A more significant advantage may be found in the fact that cilia and flagella are central elements in the morphogenesis of the three cell types discussed here. In trypanosomes, the flagellum is attached along the length of the cell body from the onset of assembly. The flagellum guides cell morphogenesis and defines the axis of cytokinesis [12, 37]. Some stages even use a flagella connector to position the new flagellum alongside the existing one [52]. In spermatozoa, the axoneme of the flagellum is fully elongated before the cascade of morphogenetic events leading the emergence of the typical elongated shape of the cell body. These occur in a precise manner and are articulated around the flagellum [9, 53]. In photoreceptors, the large outer segment develops from the cilium and is actually derived from the fusion of ectosomes that originated from it [54]. In all three situations, a stable axoneme could ensure correct morphogenesis for complete differentiation.

The grow-and-lock model raises two important questions: how to control the growth rate and the timing of the locking event? Modulating the amount of IFT trains or their loading with tubulin could impact the growth rate, as shown here for *KIN2A2B^RNAi^* cells. The amount of IFT proteins in the cell body largely exceeds that in the flagellum [13, 22] and the amount of *IFT* gene transcripts increases during flagellum assembly [55]. Similarly, a soluble pool of tubulin is available but as said above its loading on IFT trains cannot be assessed experimentally for the moment.

The timing of the locking event could be controlled at the level of the flagellum itself, which would require a way to measure flagellum length via a length sensor. This could function on the principle of the time-of-flight model that proposes that a sensing molecule traffics in association with IFT proteins and undergoes some modification in the flagellum. As the flagellum elongates, the time spent for a trip increases and the proportion of modified sensor becomes higher [56]. In induced *KIN2A2B^RNAi^* cells, the speed of IFT is reduced by ~30%, meaning that the time spent in the flagellum increases, what could trigger a premature locking event when the flagellum reaches 70% of the theoretical length. The length of the mature flagellum in induced *KIN2A2B^RNAi^* cells is compatible with this result. The grow-and-lock model could therefore function with a length sensor system, as proposed in other organisms with more dynamic flagella [57]. Alternatively, the locking event could be controlled by another cellular process that is timely regulated, such as progression through the cell cycle or through differentiation as observed here where the locking event is triggered prior cell division. The cell cycle and cilia assembly have long been known to be tightly linked [58] and a connection between the control of ciliary length and the cell cycle would make sense.

## ACKNOWLEDGEMENTS

We acknowledge Derrick Robinson (Bordeaux, France), Paul McKean (Lancaster, UK) and Robert Lee Douglas (Berkeley, USA) for generous gifts of antibodies and Linda Kohl (Paris, France) for critical reading of the manuscript. We thank Adeline Mallet for the mNG::IFT81 *KIN2A2BRNAi* cell line. We are thankful to the Ultrastructural BioImaging Plateforme for providing access to their equipment. E.B. is supported by fellowships from French National Ministry for Research and Technology (doctoral school CDV515) and from La Fondation pour la Recherche Médicale (FRM) (FDT20170436836). B.M. was supported by a Roux post-doctoral fellowship from the Institut Pasteur. This work is funded by ANR grants (11-BSV8-016 and 14-CE35-0009-01), by a French Government Investissement d’Avenir programme, Laboratoire d’Excellence “Integrative Biology of Emerging Infectious Diseases” (ANR-10-LABX-62-IBEID) and by La Fondation pour la Recherche Médicale (Equipe FRM DEQ20150734356). The funders had no role in the study design, data collection and analysis, decision to publish or preparation of the manuscript.

## AUTHOR CONTRIBUTIONS

E.B., B.M., B.R. & P.B. conceived and designed the experiments; E.B., B.M. and T.B. performed the experiments; T.B. prepared the figures, E.B. and P.B. wrote the manuscript. All authors commented on the manuscript.

## STAR METHODS

### CONTACT FOR REAGENT AND RESOURCE SHARING

Further information and requests for resources and reagents should be directed to and will be fulfilled by Philippe Bastin

### EXPERIMENTAL MODEL AND SUBJECT DETAILS

The pleomorphic strain *T. brucei* AnTat1.1E [59] was used for transformation with p2675TdTIFT81. Cells were cultured in SDM79 medium [60] supplemented with hemin, 10% fetal bovine serum and 10mM glycerol. All the other procyclic *T. brucei* cell lines were derivatives of the strain 427 and grown in SDM79 medium with hemin and 10% fetal bovine serum. All the cells were cultivated at 27°C. The 29–13 cell line expressing the T7 RNA polymerase and the tetracycline-repressor has been described previously [61]. For generation of the *KIN2A2B^RNAi^* cell line, a 489-nucleotide fragment of *KIN2A* (Tb927.11.13920) was amplified by PCR flanked by HindIII and XhoI sites and cloned in the compatible sites of the pZJM vector. The *KIN2B* (Tb 927.5.2090) fragment was generated by chemical synthesis by GeneCust Europe (Dudelange, Luxembourg). Genecust cloned these fragment into the pZJM vector [32], allowing tetracycline-inducible expression of dsRNA generating RNAi upon transfection in the 29-13 recipient cell line. The dsRNA is expressed from two tetracycline-inducible T7 promoters facing each other in the pZJM vector. Primers were selected using the RNAit algorithm to ensure that the fragment lacked significant identity to other genes to avoid cross-RNAi [62]. For generation of the *KIN2A2B^RNAi^* expressing TdT::IFT81 cell line and AnTat1.1E expressing TdT::IFT81, the first 500 nucleotides of the *IFT81* gene (Gene DB number Tb927.10.2640) were chemically synthesised (GeneCust, Luxembourg) and cloned in frame with the TdTomato gene within the HindIII and ApaI sites of the p2675 vector [63].

The construct was linearised within the *IFT81* sequence with the enzyme XcmI and nucleofected [64] in the *KIN2A2B^RNAi^* or the AnTat1.1E cell line, leading to integration by homologous recombination in the *IFT81* endogenous locus and to expression of the full-length coding sequence of IFT81 fused to TdTomato. To construct the p2675mNeonGreenIFT81 plasmid, the *mNeonGreen* gene [39] was chemically synthesised (GeneCust, Luxembourg) with HindIII and ApaI site and cloned in the corresponding site of the p2675YFPIFT81 [28] to replace the *YFP* gene. The vector was linearised with XcmI and nucleofected in the *KIN2A2B^RNAi^* cell line as described above. Transfectants were grown in media with the appropriate antibiotic concentration and clonal populations were obtained by limited dilution.

### METHOD DETAILS

#### De-induction experiments

For de-induction experiments, *KIN2A2B^RNAi^* cells were grown for 6 days in the presence of tetracycline and then washed in four times in SMD79 supplemented with serum and hemin before being returned to culture either in the presence of tetracycline (induced control) or in the absence of tetracycline (de-induced sample) during 16 hours. The experiment was reproduced 3 times.

#### Inhibition of cell division

For inhibition of cell division, teniposide (Sigma SML0609), a topoisomerase II inhibitor was dissolved in DMSO and added to trypanosome cultures at a final concentration of 200 μM [44] during 8 hours (wild-type strain, 3 independent experiments), 16 or 24 hours (*KIN2A2B^RNAi^* strain, 1 and 3 independent experiments, respectively) and. In the control flask, the same volume of DMSO was added (63 μL).

#### Scanning electron microscopy

For scanning electron microscopy, samples were fixed overnight at 4°C with 2.5% glutaraldehyde in 0.1 M cacodylate buffer (pH 7.2) and post-fixed in 1% OsO4 in 0.1 M cacodylate buffer (pH 7.2)[65]. After serial dehydration, samples were critical-point dried (Emitech K850 or Balzers Union CPD30) and coated with gold (Jeol JFC-1200 or Gatan Ion Beam Coater 681). Observations were made in a JEOL 7600F microscope.

#### Immunofluorescence and live cell imaging

Cultured parasites were washed twice in SDM79 medium without serum or in Phosphate Buffer Saline (PBS), and spread directly onto poly-L-lysine coated slides. The slides were air-dried for 10 min, fixed in methanol at −20°C for 30 s and rehydrated for 10 min in PBS. For immuno-detection, slides were incubated with primary antibodies diluted in PBS with 0.1% Bovine Serum Albumin (BSA) for 1 h at 37°C. Three washes of 10 min were performed and the secondary antibody diluted in PBS with 0.1% BSA was added to the slides. After an incubation of 45 min at 37°C, slides were washed three times in PBS for 10 min and DAPI (2 μg/μl) was added. Slides were mounted with coverslips using ProLong antifade reagent (Invitrogen). The antibodies used were the Mab25 monoclonal antibody recognising TbSAXO1, a protein found all along the trypanosome axoneme [66], an anti-IFT172 mouse monoclonal antibody diluted at 1/200 [13], an anti-FLAM8 rabbit polyclonal 1/500 (kind gift of Paul McKean, Lancaster University, UK). Subclass-specific secondary antibodies coupled to Alexa488, Alexa647 or Cy3 (1/400; Jackson ImmunoResearch Laboratories, West Grove, PA) were used for double labelling. Sample observation was performed using a DMI4000 microscope equipped with a 100X NA 1.4 lens (Leica, Wetzlar, Germany) and images captured with an ORCA-03G Hamamatsu camera. Pictures were analyzed using ImageJ 1.47g13 software (National Institutes of Health, Bethesda, MD) and images were merged and superimposed using Adobe Photoshop CC. For fluorescence quantification, we have used the Raw Integrated Density values and removed the background at all these values. For live video microscopy, cells were covered with a coverslip and observed directly with the DMI4000 microscope at room temperature. Videos were acquired using an Evolve 512 EMCCD Camera (Photometrics, Tucson, AZ), driven by the Metavue acquisition software (Molecular Probes, Sunnyvale, CA). IFT trafficking was recorded at 100 (AnTat1.1E expressing TdT::IFT81) or 250 (*KIN2A2B^RNAi^* expressing TdT::IFT81) milliseconds per frame during 30 seconds. Kymograph analysis was used to quantify anterograde and retrograde IFT. Briefly, the technique is based on an adaptive and directional band-pass filtering method that allows the separation of trails of opposite directions. The filtering method exploits the curvelet analysis of the kymograph image to automatically adapt to the trains characteristics and select oriented features. Several individual particle indicators are automatically extracted using Quia software as a function of time from the set of trajectories: the train length, its intensity averaged over the particle area, and its velocity [22, 67]. The numbers of analysed cells and of measured trains are indicated in figure or table legends. For length measurements, the Mab25 staining of the axoneme was taken as reference using ImageJ.

#### Western blot analysis

Cells were washed in PBS and boiled in Laemmli loading buffer before SDS-PAGE separation, loading 20 μg of total cell protein per lane. Proteins were transferred overnight at 25V at 4°C to polyvinylidene fluoride membranes (PVDF), then blocked with 5% skimmed milk in PBS-Tween 0.1% (PBST) and incubated with primary antibodies diluted in 1% milk and PBST. The anti-KIN2B (a kind gift of Robert L. Douglas, Berkeley)[31] serum was diluted 1/100. As loading controls, antibodies against ALBA proteins [68] diluted 1/500 were used. Three membrane washes were performed with PBST for 5 minutes. Species-specific secondary antibodies coupled to horseradish peroxidase (GE Healthcare) were diluted 1/20,000 in PBST containing 1% milk and incubated for 1 hour. Final detection was carried out using an enhanced chemoluminescence kit and a high performance chemoluminescence film according to manufacturer’s instructions (Amersham, Piscataway, NJ).

For fractionation in detergent, cells were washed twice in PBS by 5 minutes at 500 *g* and the pellet was incubated for 2 minutes in Nonidet P-40 1% in PEM buffer in the presence of protease inhibitors (Sigma P8340). After centrifugation for 2 minutes at full speed, the supernatant (soluble fraction) was separated from the pellet (cytoskeletal fraction). Cells without treatment were used as control (total extract). Samples were loaded on gel and treated as above for transfer on membranes. Tubulin was detected with the TAT-1 monoclonal antibody [69] and IFT22 that was used here as a soluble marker was detected with a mouse anti-IFT22 antiserum [70]. See also Figure S4.

### QUANTIFICATION AND STATISTICAL ANALYSIS

Statistical analyses were done with Kaleidagraph v4.5.2 using ANOVA test with Turkey HSD (α=0.5). This test was selected to detect if there were statistically significant differences between the means, knowing that variance of each mean appeared similar. Graphs were drawn using Kaleidagraph v4.5.2. All errors correspond to the standard deviation of the population. Statistically significant differences are indicated with one (p<0.001) or two stars (p<0.0001). The number of samples analysed for each experiment is indicated in figure or table legends.

## SUPPLEMENTAL INFORMATION

**Supplementary Videos**

**Video S1: Visualisation of TdT::IFT81 in AnTat1.1E cells, Related to Figure 1 and Table S1.**

TdT::IFT81 is found inside the trypanosome flagellum where it travels by IFT. Live procyclic, wild-type *T. brucei* cell transfected with TdT::IFT81 observed by time-lapse epifluorescence microscopy using a DMI4000 microscope at room temperature. Frames were taken every 100 ms for 30 s by an Evolve 512 EMCCD Camera. Example of a cell with a single flagellum.

**Video S2: Visualisation of TdT::IFT81 in AnTat1.1E cells, Related to Figure 1 and Table S1.**

Same as Video S1 but example of a cell with two flagella, the new one being at early phase of assembly.

**Video S3: Visualisation of TdT::IFT81 in 1.1E cells, Related to Figure 1 and Table S1.** Same as Video S1 but example of a cell with two flagella, the new one being at an intermediate phase of assembly.

**Video S4: Visualisation of TdT::IFT81 in AnTat1.1E cells, Related to Figure 1 and Table S1.**

Same as Video S1 but example of a cell with two flagella, the new one being at a late phase of assembly.

**Video S5: Visualisation of TdT::IFT81 in *KIN2A2B^RNAi^* cells, related to Figure S2 and Table 1.**

This is an example of a non-induced cell showing robust IFT. IFT proteins are found at the base of the flagellum and as motile trains trafficking both ways in the flagellum.

**Video S6: Visualisation of TdT::IFT81 in *KIN2A2B^RNAi^* cells, related to Figure S2 and Table 1.**

This is an example of a cell induced for 6 days where the frequency of IFT is much reduced whereas the total amount of IFT protein at the base is significantly increased.

